# A 5-day course of rTMS before pain onset ameliorates future pain and increases sensorimotor peak alpha frequency

**DOI:** 10.1101/2024.06.11.598596

**Authors:** Nahian S Chowdhury, Khandoker Taseen, Alan Chiang, Wei-Ju Chang, Samantha K Millard, David A Seminowicz, Siobhan M Schabrun

## Abstract

Repetitive transcranial magnetic stimulation (rTMS) has shown promise as an intervention for pain. An unexplored research question is whether the delivery of rTMS *prior to pain onset* might protect against a future episode of prolonged pain. The present study aimed to determine i) whether 5 consecutive days of rTMS delivered prior to experimentally-induced prolonged jaw pain could reduce future pain intensity and ii) whether any effects of rTMS on pain were mediated by changes in corticomotor excitability (CME) and/or sensorimotor peak alpha frequency (PAF). On each day from Day 0-4, forty healthy individuals received a single session of active (n = 21) or sham (n = 19) rTMS over the left primary motor cortex. PAF and CME were assessed on Day 0 (before rTMS) and Day 4 (after rTMS). Prolonged pain was induced via intramuscular injection of nerve growth factor (NGF) in the right masseter muscle after the final rTMS session. From Days 5-25, participants completed twice-daily electronic dairies including pain on chewing and yawning (primary outcomes), as well as pain during other activities (e.g. talking), functional limitation in jaw function and muscle soreness (secondary outcomes). Compared to sham, individuals who received active rTMS subsequently experienced lower pain on chewing and yawning. Although active rTMS increased PAF, the effects of rTMS on pain were not mediated by changes in PAF or CME. This study is the first to show that rTMS delivered *prior* to pain onset can protect against future pain and associated functional impairment. Thus, rTMS may hold promise as a prophylactic intervention for persistent pain.

---

Chronic pain is a global health concern, impacting patient quality of life and healthcare systems [46;53;118]. Unfortunately, current interventions (e.g. surgery, pharmacological, psychological, exercise) demonstrate at best, modest improvements in pain and function [11;98;115;116]. A critical limitation of most treatments is that they are applied once pain has already become chronic and maladaptive nervous system plasticity is entrenched [30;52]. Treatments that target features of cortical activity associated with chronic pain susceptibility *before pain begins* have potential to interrupt the transition to chronic pain and improve clinical outcomes.

High frequency (>5Hz) repetitive transcranial magnetic stimulation (rTMS) is a promising intervention for pain [43;111]. Single and multi-session 10Hz rTMS delivered to the primary motor cortex (M1) after the onset of experimental pain and in patients with chronic pain has been shown to reduce pain severity compared to sham [7;14;58;59;100]. Despite this, little research has investigated the prophylactic effect of rTMS applied prior to pain onset on future pain severity. Several studies demonstrated that a single session of high frequency rTMS [9;61;69;70;117] delivered within an hour before transient pain led to reduced pain sensitivity and severity in pain-free individuals. However, no study has explored these effects using a prolonged pain model, such as pain induced by intramuscular injections of nerve growth factor (NGF) [45;89]. This is a critical question, as interventions that can ameliorate future prolonged pain might also be effective in hindering the transition from acute to chronic pain. NGF injections induce prolonged musculoskeletal pain [17] that mimics the duration, time course, functional limitation and hyperalgesia associated with chronic pain conditions [8;89]. This makes the NGF model a highly standardised and clinically relevant model for exploring the prophylactic effects of rTMS on a future prolonged pain episode.

The prophylactic effects of rTMS on pain might be mediated by two features of cortical activity shown to predict an individual’s pain sensitivity - peak alpha frequency (PAF), which refers to the dominant oscillatory frequency in the 8-12Hz range [16] and corticomotor excitability (CME), which is indexed via the magnitude of motor responses to single-pulse TMS to M1 [112]. Several studies have shown that slower sensorimotor PAF and lower CME predicts higher future pain severity [32-34;49;50]. This would suggest that interventions that can increase PAF and CME might protect against future pain. 10Hz rTMS to M1 is thought to directly increase the excitability of motor cortical neurons through long-term potentiation mechanisms [3;24], or via entrainment of intrinsic alpha oscillators as a result of synchronization with external stimulation [74;106]. Consistent with these explanations, studies have reported increases in CME and PAF following single and multi-session high frequency rTMS [2;29;64;67;73]. Thus, it is plausible that the prophylactic effects of rTMS on pain is mediated by increases in PAF and CME.

The present study aimed to determine whether 5 consecutive days of rTMS delivered *prior* to the onset of NGF-induced temporomandibular pain could subsequently reduce pain severity compared to sham rTMS. A secondary aim was to determine whether the prophylactic effects of rTMS were mediated by changes in PAF and CME. It was hypothesised that active rTMS would lead to lower pain severity compared to sham, and that reductions in pain would be mediated by increases in PAF and CME.

## Methods

### Design

This study used a longitudinal, randomised, sham-controlled, parallel design to follow healthy individuals over the course 26 days. Figure 1 shows the experimental protocol. Participants attended five lab sessions (Day 0-4). On the Day 0 visit, PAF and CME were assessed, followed by delivery of either active or sham rTMS. On Days 1, 2 and 3, participants received the rTMS intervention only. On Day 4, participants received the final session of rTMS, followed immediately by a repeat assessment of PAF and CME. Intramuscular injection of NGF was given into the right masseter muscle at the end of the Day 4 session to induce prolonged pain and dysfunction mimicking temporomandibular disorder (TMD) [89]. From Days 5-25, participants completed twice daily electronic dairies comprising measures of pain and function. Written, informed consent was obtained prior to study commencement. The study was approved by the UNSW ethics committee (HREC reference number: HC220602). Data collection was conducted at Neuroscience Research Australia (NeuRA).

**Figure 1.**
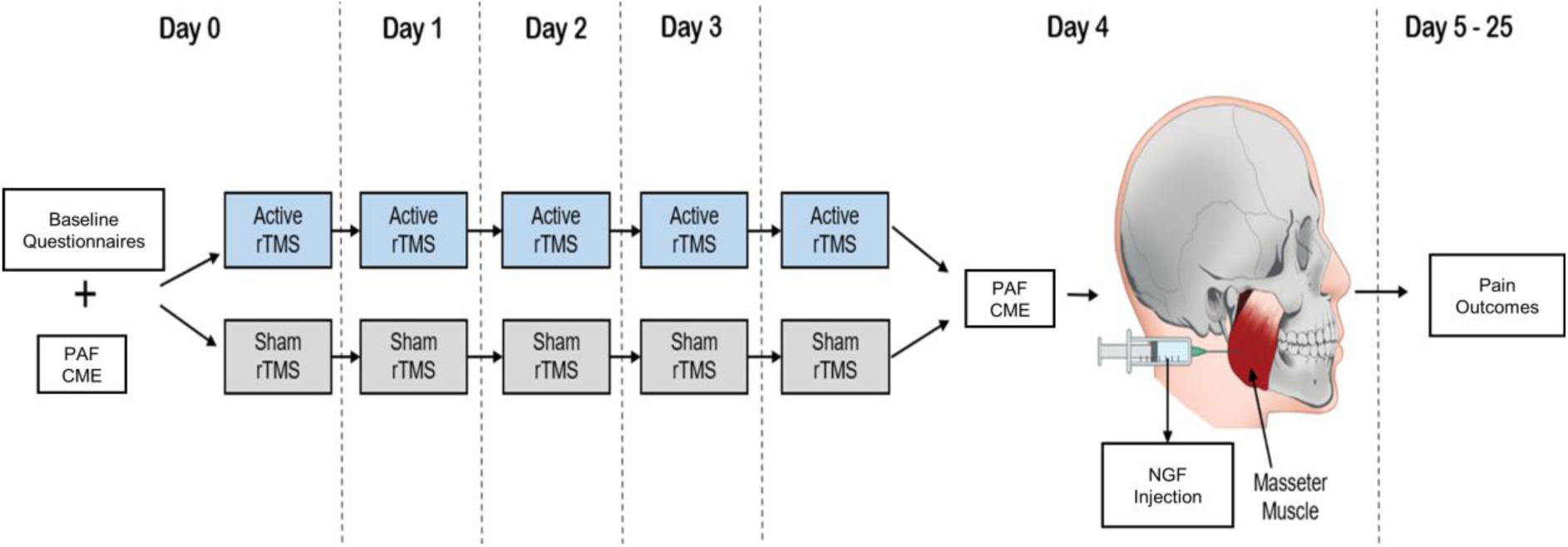
Diagram of the experimental protocol

### Participants

Participants were recruited through notices posted online, at universities across Sydney, Australia or by contacting participants held on a database at NeuRA. Forty-one healthy individuals (23 female and 18 male; age 23 ± 4 [mean ±SD]; range 18-35) were recruited. Participants were included if they were aged between 18-44. This was justified based on data from the OPPERA prospective cohort study that demonstrated an incidence rate of first onset TMD of 2.5% per annum among 18 to 24-year olds and 4.5% per annum among 35 to 44-year olds [99]. Participants were excluded if they presented with any acute pain, had a previous history of chronic pain, major medical complaints, psychiatric or salivary gland conditions, were pregnant and/or lactating, or were contraindicated for TMS (e.g., metal implants in the skull, epilepsy) as assessed using the Transcranial Magnetic Stimulation Adult Safety Screen Questionnaire [85]. A sample size calculation was conducted (G*Power 3.1.9.7) based on effect sizes, which varied between 1.08 and 2.17, from previous studies examining the influence of active vs. sham rTMS on NGF-induced pain[13;26], producing a sample size between 5-15 per group to achieve 80% power at .05 significance. Given that no previous study has applied rTMS before prolonged pain onset, we conservatively selected a sample size of 20 individuals in each group.

### Randomization/Blinding Procedure

Prior to the study, participants were randomly assigned to either the active or sham condition using a random number list generator (https://www.random.org/lists/). The allocations were revealed to the experimenter (KT, AC or WC) delivering rTMS on Day 0 via an automated email notification. The allocations were not known by the experimenter (NC) that collected PAF and CME data, administered the injection of NGF, performed data pre-processing and prepared the analysis plan. Group allocations were revealed to experimenter NC when the data collection, pre-processing and analysis plan was complete. Participants were not informed of the existence of two conditions at any time throughout the study (Day 0-25). Participants were debriefed (experimenter KT) after completing the study (contacted by phone or email) to explain that they were randomly assigned to either an active or sham intervention and were asked which condition they believed they received. Subsequently, participants’ group allocation was notified by a follow-up email.

### Data Collection Procedures

#### Baseline Questionnaires

On Day 0, prior to rTMS, participants completed a series of questionnaires including the (1) Pain Catastrophizing Scale [101], (2) Perceived Stress Scale [104], (3) Sleep Scale, (4) Patient health questionnaire [57], (5) Pennebaker Inventory of Limbic Languidness Questionnaire [78], (6) Short-Form-8 Health Questionnaire [109] and (7) The Brief Pain Inventory Pain Severity and 7-item Interference subscales [55;103]. These questionnaires would help characterize the physical and mental health profile of the active and sham groups.

#### Peak Alpha Frequency

Resting state EEG was collected in line with previous methods [33]. Participants were seated in a comfortable chair. Scalp EEG was recorded using the Brain Products platform (BrainVision Recorder, Vers. 1.22.0101 with actiCHamp Plus, Brain Products GmbH, Gilching, Germany). Recording was done at a sampling rate of 5000 Hz. Signals were recorded from 63 active electrodes (actiCAP slim, Brain Products GmbH, Germany), embedded in an elastic cap (EASYCAP, EASYCAP GmbH) in line with the 10–10 system. Recordings were referenced online to ‘FCz’ and the ground electrode placed on ‘Fpz’. Electrode impedances were maintained below 25 kOhms. Based on previous research [28;54] and the actiCHamp manual, we anticipated negligible signal loss with this impedance cut-off. Once setup was complete, the lights were switched off with ambient noise reduced to a minimum. Participants were instructed to relax and keep their eyes closed while remaining awake. The resting-state EEG signal was then recorded for 5 minutes.

#### Corticomotor Excitability

TMS was used to map the corticomotor representation of the right masseter muscle, in line with previously described techniques [20;88;90]. Participants wore a swim cap with a grid of 1cm x 1cm resolution. Single-pulse, monophasic stimuli were delivered to the left hemisphere using a Magstim stimulator and a figure-of-eight coil. Bipolar surface electrodes were used to record electromyographic (EMG) activity from the masseter muscle, with active (muscle belly) and reference (tendon) electrodes placed along the mandibular angle, and the ground electrode was placed on the right acromion process. EMG signals were amplified (x2000), with a high pass (16Hz) and low pass (1000 Hz) filter applied, and digitally sampled at 5kHz.

It is common to assess CME from the masseter muscle during active contraction that is at a certain percentage of the maximum voluntary contraction (MVC) [23;83], as it is challenging to obtain reliable MEPs from the masseter muscle at rest [10;113]. Participants were instructed to contract at 20% MVC which provided a balance between obtaining sufficiently large MEPs for analyses [65] and minimizing fatigue. To measure MVC, participants were verbally encouraged to clench their jaw as hard as possible for 3 seconds on 3 separate trials, with a 1-minute rest break in between trials. The MVC was computed based on the average of the 3 trials, and 20% MVC was computed based on this average. Participants practiced maintaining a masseter contraction of 20% MVC by receiving live visual feedback from the EMG signal prior to TMS recording.

Consistent with previous studies investigating optimal coil orientation for inducing masseter MEPs, an angle of 90 degrees between the anterior-posterior line and the coil handle was used [39]. This orientation induced current in the lateral-to-medial direction. The scalp site evoking the largest MEP (i.e., the “hotspot”) during active contraction was then determined. The active motor threshold (AMT) was determined using the ML-PEST (maximum likelihood strategy using parametric estimation by sequential testing) algorithm [6] which estimates the minimum TMS intensity required to induce a reliable MEP with a 50% probability . A reliable MEP was identified if the EMG waveform between ∼5-15ms [63;77] after the TMS pulse was visually larger in amplitude relative to background contraction. The TMS intensity for mapping was set at 120% AMT.

The procedure for mapping has been described in detail previously [87;90]. While participants maintained contraction of the masseter muscle, three stimuli were delivered at each location around the grid, starting at the hotspot. The number of stimulation sites was pseudorandomly increased until an MEP was no longer observed (i.e., no reliable MEP in all 3 trials at all border sites). In line with the approach used in previous studies [25;95;102], the hotspot location and test stimulus intensity determined on Day 0 was used for subsequent mapping procedures on Day 4. This approach ensures that any changes in motor cortical maps occurring as a result of rTMS can be observed. To confirm that the AMT did not change, we re-assessed AMT on Day 4.

#### rTMS

We employed a rTMS protocol used in previous studies investigating the analgesic effects of rTMS [13;93]. High-frequency 10Hz, biphasic active rTMS was applied over the left primary motor cortex using a Magstim Super Rapid2 stimulator and a figure-of-eight coil. The location and intensity of rTMS was calibrated based on resting motor threshold (RMT) of the first dorsal interosseous (FDI) muscle rather than the targeted masseter muscle. This is due to the difficulty involved in obtaining reliable masseter responses to TMS at rest [10;113] and given previous studies have demonstrated non-somatotopic effects of rTMS on pain [4;51;56]. Surface bipolar EMG electrodes were placed over the FDI to record MEPs. The coil was orientated at 45° to the midline. The motor hotspot for the FDI was determined, which corresponded to the scalp site which generated the largest MEP at the targeted FDI. The RMT was determined using ML-PEST [6], which estimated the minimum TMS intensity required to induce a MEP of 50 microvolts with a 50% probability. This was reassessed at each session to account for potential changes in CME in response to rTMS [76;94]. During rTMS, the coil was held at 90° to the midline [60]. 3000 stimuli (10 Hz, 30 trains of 10 seconds, 20-second intertrain interval) were delivered at 90% of RMT. For sham rTMS, a sham coil that looked identical to the active rTMS coil but produced only audible clicks was used to deliver the stimulation protocol identical to the one used for active rTMS.

#### NGF Injection

The NGF injections were provided at the end of the Day 4 session. To prepare the masseter muscle for injection, the surface was cleaned using alcohol wipes. A sterile solution of recombinant human NGF (dose of 5 μg [0.2 ml]) was then administered as a bolus injection into the muscle belly of the right masseter using a 1-ml syringe with a disposable needle (27-G). The needle was inserted perpendicular to the masseter body until bony contact was reached, retracted ∼2 mm, and NGF injected. Any adverse events (e.g. excessive bleeding, swelling) were monitored.

#### Electronic Pain dairies

Questionnaires assessing jaw pain, muscle soreness, and functional limitation were completed on Day 0 and Day 4 (before and after rTMS) prior to the NGF injection, twice daily from Day 5 to 25 (after NGF injection) via automated links to an electronic dairy sent to participants at 10am and 7pm each day.

##### Primary outcomes

Pain upon functional jaw movement is a key criterion for the diagnosis of TMD [91]. Moreover, previous research has shown that, after an NGF injection to the masseter muscle, pain during chewing and yawning activities are higher compared to other activities [23;89]. As such, our primary outcomes were pain upon chewing and yawning. Pain intensity was rated on an 11-point numerical rating scale (NRS). The scale ranged from 0 = ‘no pain’ to 10 = ‘worst pain imaginable’.

##### Secondary outcomes

At each timepoint, participants also rated pain at rest, swallowing, drinking, talking, and smiling. They also completed a functional limitation scale consisting of 20 activities to assess jaw functional impairment experienced in the previous 24 hours [72]. For each item, the participant rated the ease at which they could perform the particular activity on an 11-point NRS, 0 = ‘no limitation’ and 10 = ‘severe limitation’. If an activity was not applicable, a rating was not provided or given as ‘NA’. The mean score was calculated for each participant across 3 subscales. Items 1-6 represented mastication, items 7-10 jaw mobility, and items 11-20 verbal and emotional communication. Lastly participants completed a muscle soreness scale on a modified 7-point Likert scale [45]: 0 = a complete absence of soreness, 1 = a light soreness in the muscle felt only when touched/vague ache, 2 = a moderate soreness felt only when touched/a slight persistent ache, 3 = a light muscle soreness when opening the mouth fully or chewing hard food, 4 = a light muscle soreness, stiffness or weakness when opening the mouth or chewing soft food, 5 = a moderate muscle soreness, stiffness or weakness when opening the mouth or chewing soft food, and 6 = a severe muscle soreness, stiffness or weakness that limits the ability to open the mouth or chew.

### Data Pre-processing

#### Sensorimotor Peak Alpha Frequency

EEG data processing was conducted using custom MATLAB (R2020b, The Mathworks, USA) scripts implementing the EEGLAB (eeglab2019_1) [27] and Fieldtrip (v20200215) toolboxes [75]. The pre-processing pipeline matched those used in previous studies investigating the relationship between PAF and pain [20;32]: data downsampled to 500 Hz, re-referenced to common average, and band-pass filtered between 2 and 100 Hz using an FIR filter. Channel data was visually inspected, and overtly noisy channels were removed, and the signal re-referenced. Data were segmented into 5s epochs, and epochs containing marked muscular artefacts were manually rejected. Independent component analysis (runica in fieldtrip) was applied to the data. We visually inspected the frequency-spectra of the components, and identified components that had a clear alpha peak (8–12 Hz) and a scalp topography suggestive of a source predominately over the sensorimotor cortices [32]. The power spectral density of the chosen sensorimotor component was derived in 0.2 Hz bins. The 2–50 Hz range was extracted using Fast Fourier Transform. A Hanning taper was applied to the data to reduce edge artefacts. Sensorimotor alpha power was calculated as the sum of all z-transformed spectral amplitudes in the 8-12Hz window. Sensorimotor PAF was calculated using the Centre of Gravity method, using the following equations:

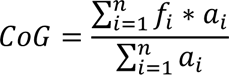

Where f_i_ is the ith frequency bin within 8-12Hz window, and a_i_ represents the spectral amplitude at f_i_.

#### Corticomotor Excitability

TMS data was processed using a custom MATLAB script. The MEP onsets and offsets were fixed to 7.3 and 15.3 ms after the TMS pulse based on our previous study [20] which showed that using a fixed MEP window produce similar map outcomes to manually choosing the offsets and onsets. The root mean square (RMS) of background EMG was determined using a fixed window between 55 and 5ms before the TMS pulse [86]. Background RMS was subtracted from the RMS of the MEP window to determine MEP amplitude. The mean MEP amplitude at each stimulation site was determined and normalized (as percentage of the maximum site) [71]. CME was indexed as map volume, which was calculated by adding the MEP amplitudes of all active sites. A stimulation was deemed active if the MEP amplitude was >10% of the maximum MEP amplitude [107].

#### Electronic Pain Diaries

We consolidated the morning and afternoon recordings into a single table to maintain a chronological order of pain intensity records, spanning from Day 5 through Day 25.

### Statistical Analyses

Means and Standard Deviations (SD) in pain outcomes, sensorimotor PAF and CME were calculated for each group. Assumptions were assessed using Shapiro-Wilk Test (for normality of pain, Sensorimotor PAF and CME data) and Mauchley’s test (for equality of variance between timepoints). Any normality assumption violations were addressed by square-root transforming the data or using non-parametric tests where applicable. Statistical significance was set to .05.

#### Effect of rTMS on pain outcomes

To determine the effect of active rTMS relative to sham rTMS, a mixed-model ANOVA was conducted with factors time (Day 5-25) and group (active vs. sham) and primary or secondary pain outcomes as the dependent measure. The group main effect assessed whether, averaged across all days (Day 5-25), active and sham groups differed on the dependent measures. The interaction effect assessed whether group level effects differed across time.

#### Effect of rTMS on Sensorimotor PAF and CME

A mixed-model ANOVA with factors: group (active vs. sham) and time (Day 0 vs. Day 4), was used to determine whether active rTMS modulated CME and sensorimotor PAF compared with sham rTMS (i.e., whether there was a group x time interaction). A mixed-model ANOVA with factors: group (active vs. sham) and time (Day 0 vs. Day 4), was also conducted to determine whether active rTMS modulated alpha power (quantified as the sum of the power from all frequency bins between 8-12Hz) compared to sham rTMS.

#### Mediation Analysis

Mediation analysis was conducted in R (version 4.2.1) using the lavaan package [84]. For the primary pain outcomes, we determined whether the effects of rTMS on pain (mean pain of first week [Days 5-11]) were mediated by changes in PAF or CME (Day 4 – Day 0) using two separate mediation models. Figure 2 shows the mediation models used, which assessed the direct effect of rTMS on PAF or CME (*a* path), the effect of change in PAF/CME on pain (*b* path), the direct effect of rTMS on pain outcome (*c’* path), and the indirect effect of rTMS on pain (*a* + *b* path). A significant indirect effect suggests the effect of rTMS on pain was mediated by changes in PAF or CME.

**Figure 2.**
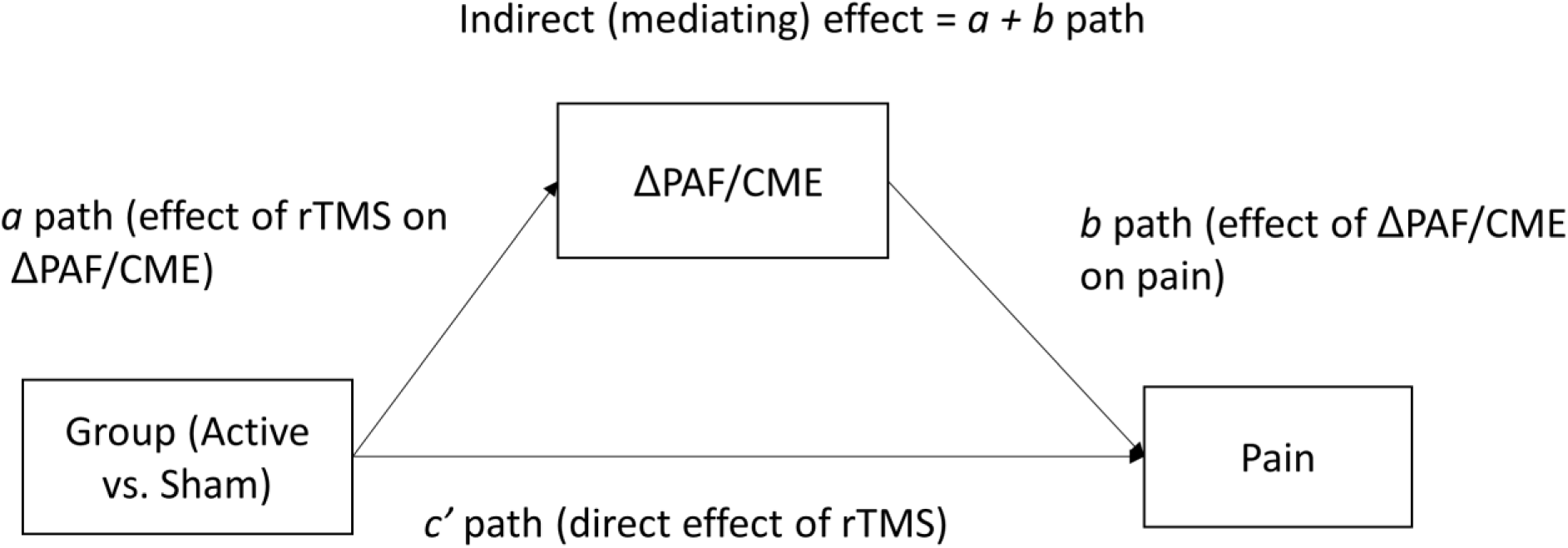
Model for the mediation analysis, which assessed the direct effect of rTMS on PAF or CME (a path), the direct effect of rTMS on pain outcome (c path), and the indirect effect of rTMS on pain (a + b path).

#### Relationship between PAF/CME prior to injection and subsequent pain intensity

To test the theoretical rationale that slower PAF/lower CME predicts higher future pain intensity, we conducted a correlation analysis to determine whether slower sensorimotor PAF and lower CME at Day 0 and Day 4 (regardless of group allocation) was associated with and average pain upon yawning and chewing of the first week (Day 5 to Day 11).

## Results

All participants completed the 5 sessions as planned, with the exception of one participant who withdrew on Day 2 due to an unwillingness to receive the subsequent NGF injection, leaving 40 participants in the final sample (21 active, 19 sham). Demographics and questionnaire data are summarised in Table 1. No adverse events were reported. Following CONSORT recommendations [68], baseline statistics comparing participant characteristics were not reported. Two participants could not be contacted at the conclusion of the study for their group allocation. For the remaining 38 participants, the proportion of participants who guessed the correct condition was not significant for both the active group (62%, *ꭓ*^2^ = 1.19, *p* = .27) and sham group (53%, *ꭓ*^2^ = .05, *p* = .82), indicating that participant blinding was successful.

**Table 1.** Participant Characteristics. Data are means (SD) or numbers (%). PCS, Pain Catastrophizing Scale; PSS, Perceived Stress Scale; PHQ, Patient health questionnaire; PILL, Pennebaker Inventory of Limbic Languidness Questionnaire; SF-8 (PCS, MCS), Short-Form-8 health questionnaire (physical component summary, mental component summary); BPI, Brief Pain Inventory Pain Severity and 7-item Interference subscales.

|  |  | **Active rTMS** | **Sham rTMS** |
| --- | --- | --- | --- |
| **Sample size** |  | 21 | 19 |
| **Sex (F,M)** |  | 11, 10 | 11, 8 |
| **Age (years)** | | 22.2 ± 3.0 | 24.4 ± 5.4 |
| **PCS** | | 8.6 ± 8.4 | 10.6 ± 8.4 |
| **PSS** | | 17.5 ± 6.6 | 18.7 ± 3.1 |
| **Sleep Scale** | **Total sleep (hours)** | 7.3 ± 1.2 | 6.9 ± 0.9 |
| | **Total sleep score** | 40.5 ± 8.3 | 43.9 ± 5.0 |
| **PHQ** | | 0.9 ± 1.4 | 0.5 ± 0.8 |
| **PILL** | | 94.7 ± 31.4 | 85.8 ± 20.8 |
| **SF-8 PCS** | | 1.9 ± 0.5 | 1.8 ± 0.5 |
| **SF-8 MCS** | | 1.9 ± 0.9 | 1.8 ± 0.7 |
| **BPI Inventory** | | 0.5 ± 1.0 | 0.1 ± 0.2 |
| **BPI Interference** | | 1.0 ± 1.3 | 0.5 ± 0.6 |
| **Correctly identified group allocation** |  | 13/21 (62%) | 9/17 (53%) |

### rTMS did not alter RMT

For the active and sham groups respectively, the mean resting motor threshold was 51.4 ± 10.9 and 59.9 ± 9.9 on Day 0, and 53 ± 11.1 and 56.4 ± 13.4 on Day 4. A two-way ANOVA (time main effect: Day 0-4, group: active vs. sham) showed resting motor threshold did not differ across Days 0-4, *F*(4, 152) = .61 *p* = .66, or between groups (group main effect: *F*(1,38) = 2.59, *p* = 0.12, group x time interaction: *F*(4,152) =1.67, *p* = .16).

### Characteristics of the NGF model

Diary completion rates were ∼90% on average (95% for the first week after injection).

We used linear interpolation to impute missing pain data, as has been done previously in situations where collection of pain outcomes yields missing data [31;108;110]. Figure 3 shows scores for the active and sham rTMS groups on the primary pain outcomes. For both groups, pain upon chewing and yawning peaked the day after the injection, with 2.2 ± 1.4 and 2.6 ± 1.7 points out of 10 respectively in the active group, and 3.1 ± 2.4 and 4.2 ± 2.4 points out of 10 in the sham group for the 10AM timepoint. Figure 4 shows the scores for other pain outcomes, functional limitation and muscle soreness across time. For both groups, functional limitation during mastication, jaw mobility and verbal communication peaked the day after the injection, with 2.5 ± 2.5, 2.1 ± 2.5 and 1.0 ± 1.5 points out of 10 respectively in the active group and 2.5 ± 2.2, 2.3 ± 1.7 and 1.2 ± 1.5 points out of 10 in the sham group for the 10AM timepoint. For both groups, muscle soreness peaked three days after the injection (Day 7), with 2.6 ± 1.6 points out of 7 in the active group and 2.6 ± 1.6 points out of 7 in the sham group, for the 10AM timepoint.

**Figure 3.**
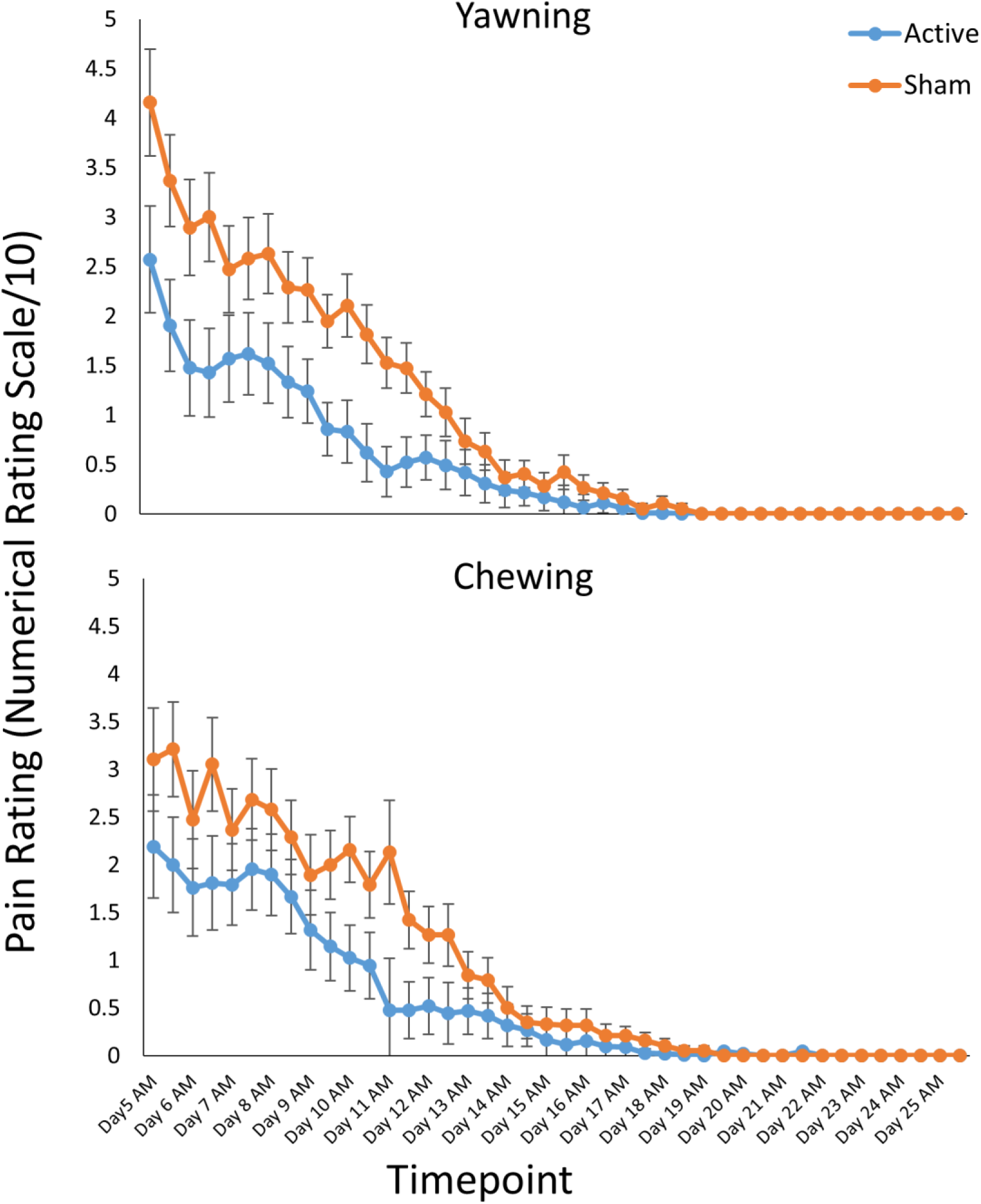
Mean ratings (±SEM) for the primary outcomes (pain upon chewing and yawning) in each group as a function of timepoint.

**Figure 4.**
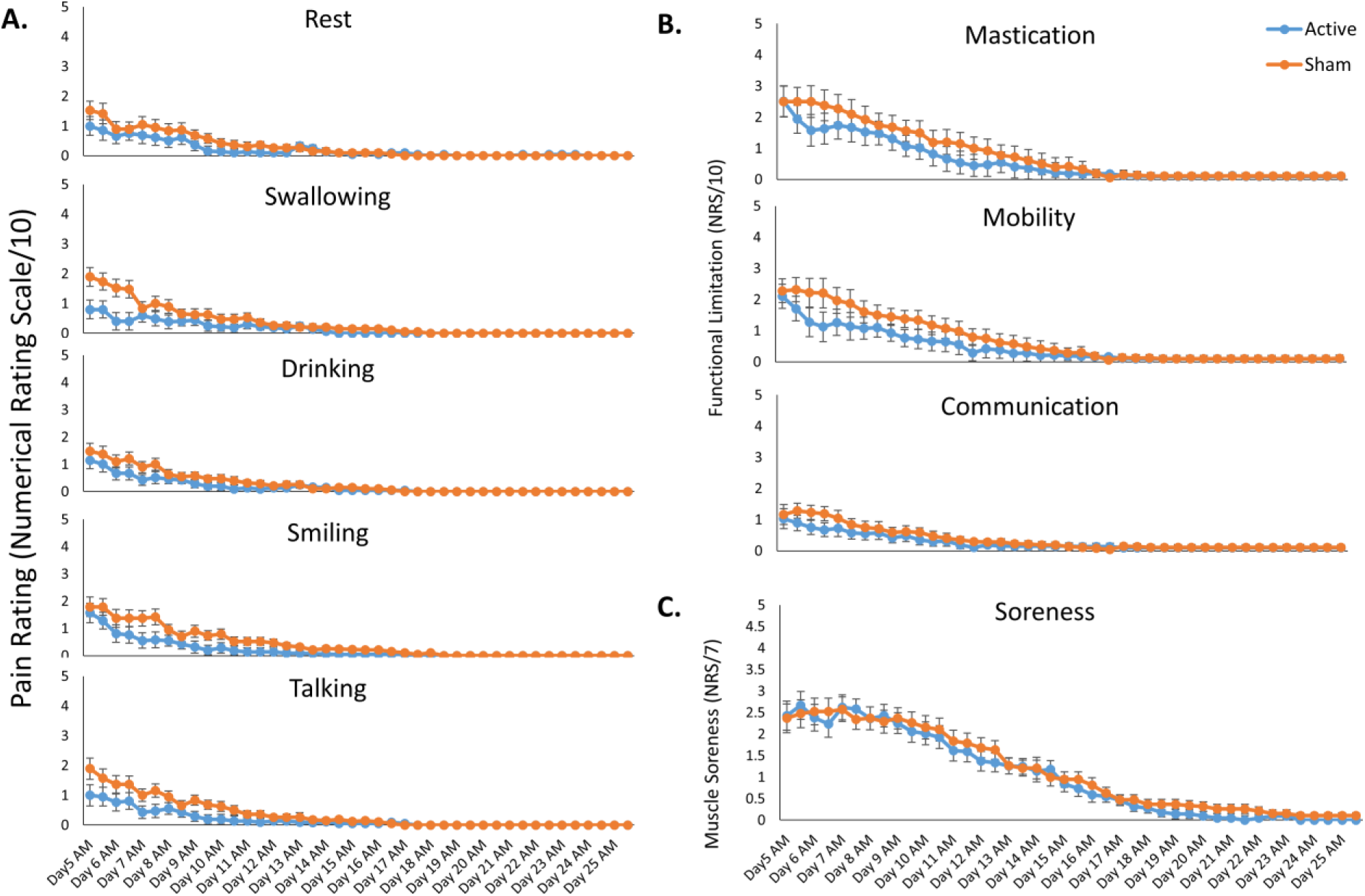
Mean ratings (±SEM) for the secondary outcomes, including pain during activities not directly relevant to the pain model (A), functional limitation (B), muscle soreness and (C) in each group as a function of timepoint.

### rTMS applied before pain onset reduces future pain severity

#### Primary outcomes

Compared to sham, those who received active rTMS experienced lower pain upon chewing and yawning, with evidence that these effects were larger at earlier timepoints (pain on chewing: time main effect, *F*(41, 1558) = 81.12, *p* < .001, group main effect, *F*(1, 38) = 5.24, *p* = .028, group x time interaction, *F*(41,1558) = 3.46, *p* < .001, group x time interaction linear trend: *F*(1, 38) = 6.17, *p* = .017; pain on yawning: time main effect, *F*(41,1558) = 74.47, *p* < .001, group main effect, *F*(41,1558) = 4.17, *p* < .001, group x time interaction, *F*(41,1558) = 4.17, *p* < .001, group x time interaction linear trend: *F*(1, 38) = 8.33, *p* = .006).

#### Secondary Outcomes

While active rTMS did not lead to lower pain upon talking or lesser functional limitation of jaw mobility when averaged across the 3 week period, stronger reductions in pain, and less functional limitation were present compared to sham at earlier timepoints (pain upon talking: time main effect, *F* (41,1558) = 21.83, *p* < .001, group main effect: *F* (1,38) = 2.20, *p* = .15, group x time interaction: *F*(41, 1558) = 1.73, *p* = .003; functional jaw mobility: time main effect, *F* (41,1558) = 55.96, *p* = .001, group main effect*, F*(1,38) = 1.14, *p* = .29, group x time interaction, *F*(41, 1558) = 2.43, *p* < .001). The rTMS intervention did not alter the severity or time course of pain associated with swallowing, smiling, drinking, or pain at rest, functional outcomes of mastication and communication, or muscle soreness (*p*’s > .05 for all group and group x time interaction effects, *p*’s < .05 for all time main effects).

### Five days of rTMS increases PAF, but does not alter CME

Figure 5 shows the mean spectral and topographical data, and violin plots of individual data of the identified sensorimotor alpha component for each rTMS group at Day 0 and Day 4. Figure 6 shows the mean masseter motor maps for each rTMS group at Day 0 and Day 4. 5 days of active rTMS modulated PAF when compared to sham rTMS (time main effect: *F*(1, 38) = .46, *p* = .50, group main effect: *F*(1, 38) = .74, *p* = .40, group x time interaction: *F*(1, 38) = 6.41, *p* = .016), with an increase in PAF on Day 4 vs. Day 0 following active rTMS (*t*_20_ = 2.34, *p* = .029). While observation of Figure 5 might suggest the increase in PAF was accounted for by an increase in power in 10-12Hz alpha band, there were no significant effects of active rTMS on alpha power when compared to sham (time main effect: *F*(1, 38) = .03, *p* = .88, group main effect: *F*(1, 38) = 1.05, *p* = .31, group x time interaction: *F*(1, 38) = 1.52, *p* = .23). rTMS did not alter CME (time main effect: *F*(1, 38) = .06, *p* = .82, group main effect: *F*(1, 38) = 1.90, *p* = .18, group x time interaction: *F*(1, 38) = 2.62, *p* = .11).

**Figure 5.**
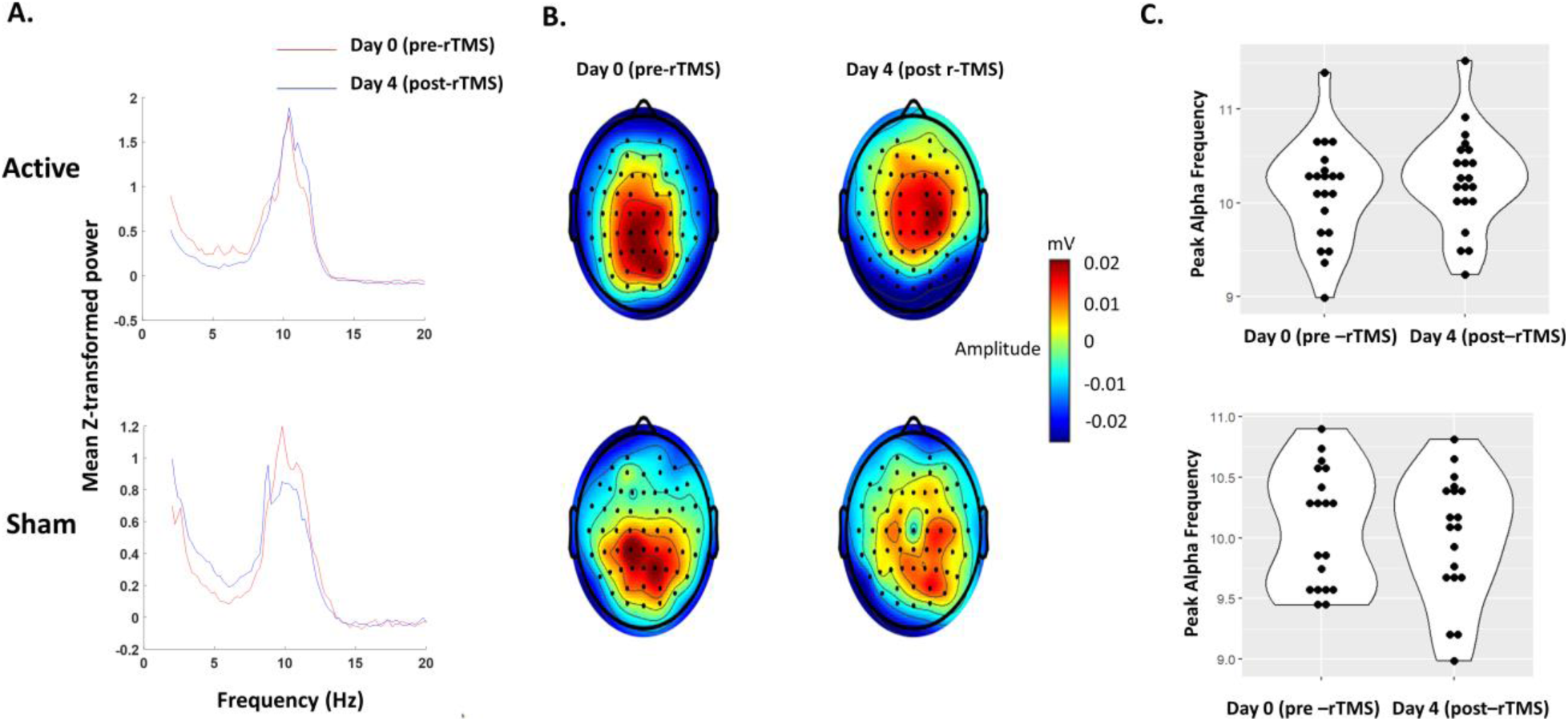
A: Mean spectral plots for each group at Day 0 (pre-rTMS) and Day 4 (post rTMS). B: Mean topographical plots of the identified sensorimotor alpha component for group at Day 0 (pre-rTMS) and Day 4 (post rTMS). C: Violin plots showing individual data points for peak alpha frequency for each group at Day 0 (pre-rTMS) and Day 4 (post rTMS)

**Figure 6.**
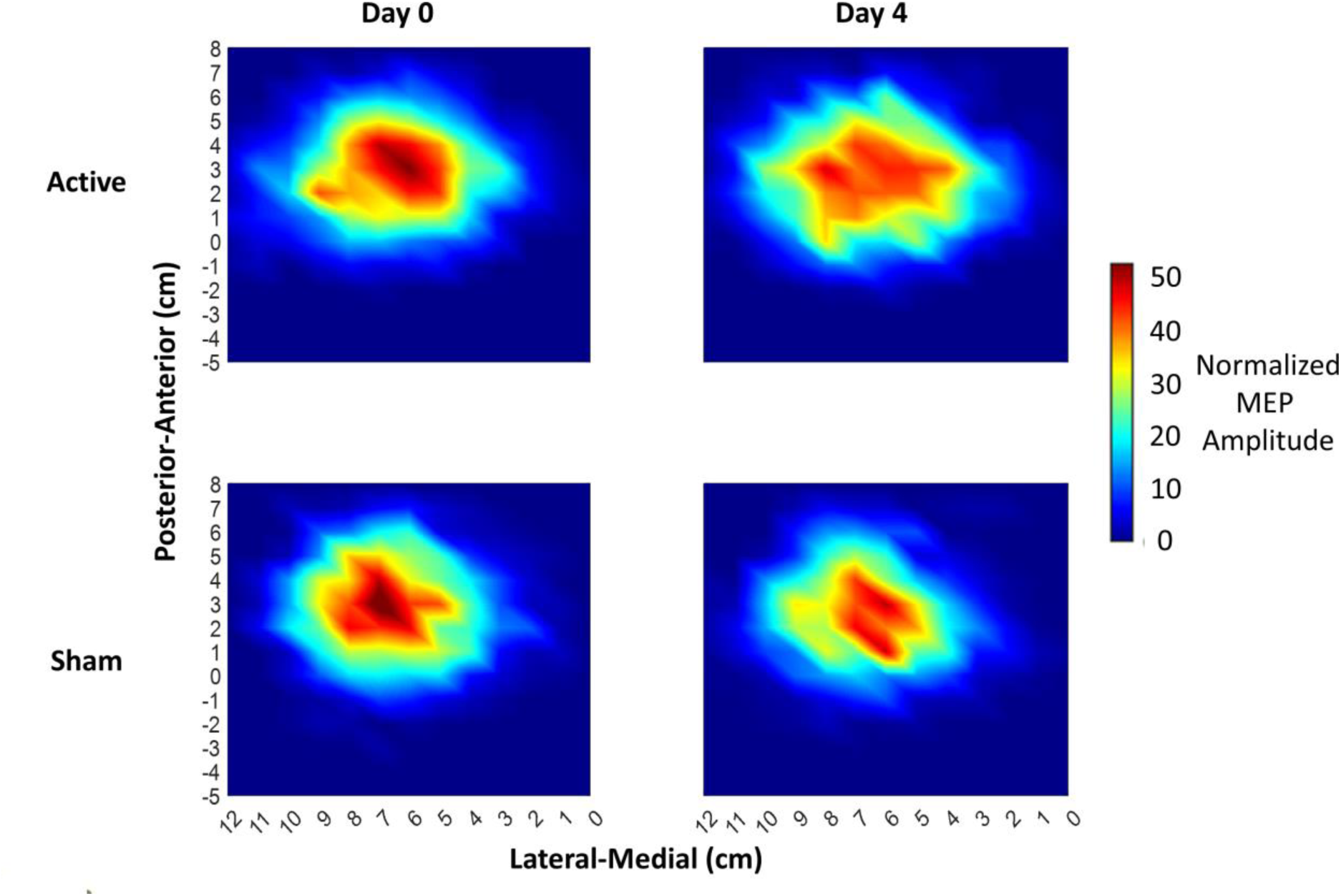
Mean masseter motor maps for each group at Day 0 (pre-rTMS) and Day 4 (post rTMS). The maps show the normalized MEP amplitude at each stimulation site relative to the vertex in both lateral-medial and posterior-anterior dimensions.

### The effects of rTMS on pain were not mediated by changes in PAF or CME

While the mediation models showed significant effects of rTMS on pain upon chewing and yawning and an effect of rTMS on PAF, there was no evidence that changes in PAF mediated the effect of rTMS on pain upon chewing (total effect: *b* = -.40, *p* = .006, direct effect of rTMS on pain: *b* = -.36, *p* = .023, effect of rTMS on PAF: *b* = .21, *p* = .009, indirect effect: *b* = -.05, *p* = .45) or yawning (total effect: *b* = -.56, *p* = .001, direct effect of rTMS on pain: *b* = -54, *p* = .004, effect of rTMS on PAF: *b* = -.40, *p* = .006, indirect effect: *b* = -.01, *p* = .86). There was also no evidence that changes in CME mediated the effect of rTMS on pain upon chewing (total effect: *b* = -.41, *p* = .006, direct effect of rTMS on pain: *b* = -.39, *p* = .01, effect of rTMS on CME: *b* = .27, *p* = .097, indirect effect: *b* = - .02, *p* = .67) or yawning (total effect: *b* = -.56, *p* = .001, direct effect of rTMS on pain: *b* = -0.52, *p* = 003, effect of rTMS on CME: *b* = .27, *p* = .097, indirect effect: *b* = -.03, *p* = .54).

### PAF and CME predict future pain severity

Slower PAF on Day 4 was associated with higher pain upon chewing (spearman’s *ρ* = -0.40, *p* = .01) and yawning (*ρ* = -0.32, *p* = .05) averaged across the first week (Days 5-11), as shown in Figure 7. Moreover, lower CME on Day 4 was associated with higher pain on yawning (*ρ* = -0.40, *p* = .01) but not chewing (*ρ* = -0.21, p = .18). There was no relationship between Day 0 CME and chewing (*ρ* = -0.04, *p* = .8) and yawning (*ρ* = -.12, *p* = .45). There was also no relationship between Day 0 PAF and pain during chewing (*ρ* = -0.24, *p* =0.14) and yawning (*ρ* = -0.27, *p* =.10).

**Figure 7.**
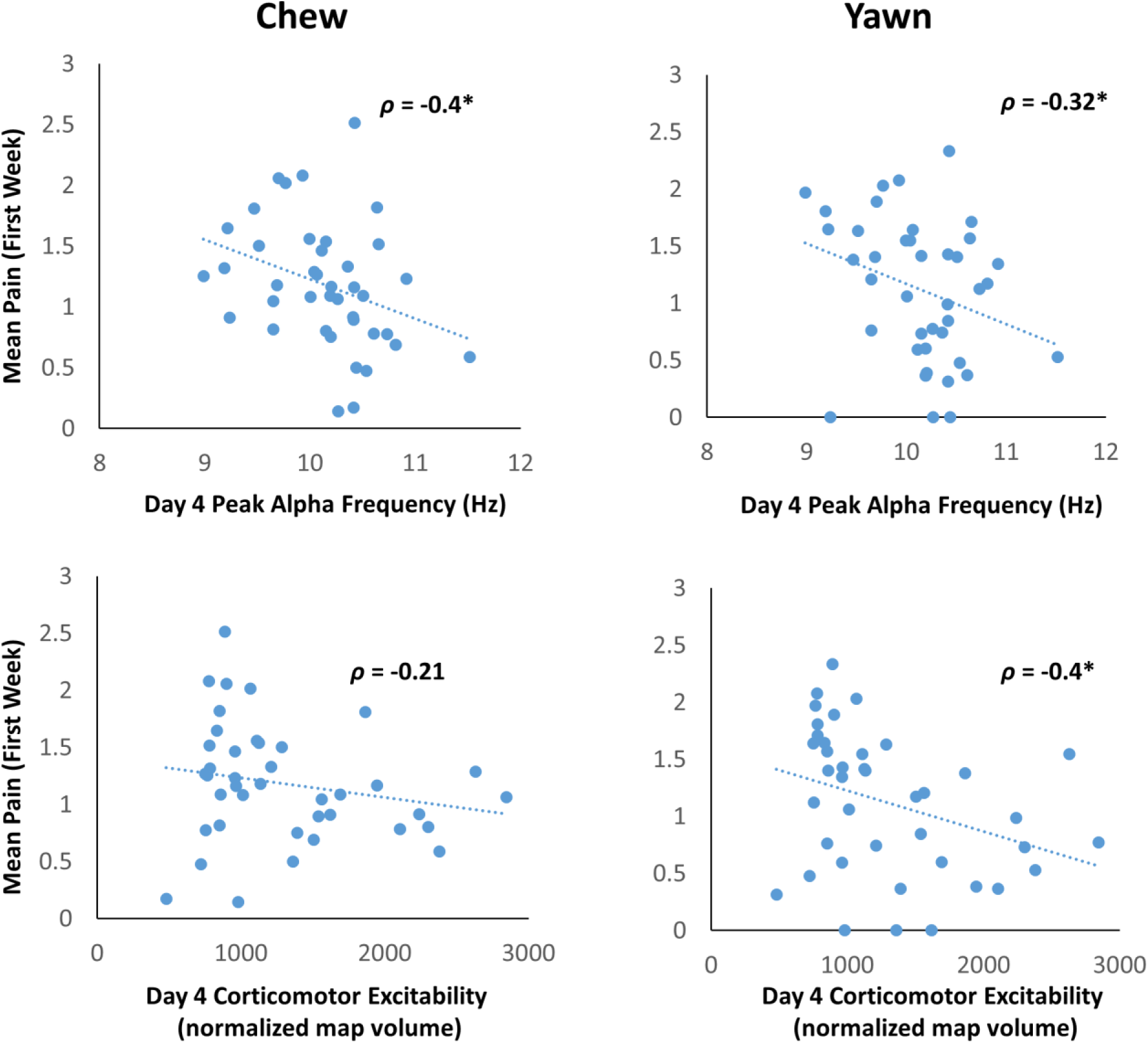
Scatter plots showing the association between peak alpha frequency and corticomotor excitability on Day 4 against the mean pain rating (chewing and yawning) across the first week (Day 5-11).

## Discussion

The present study investigated whether 5 consecutive days of rTMS over left M1 prior to the onset of prolonged temporomandibular pain could reduce future pain severity. A secondary aim was to determine whether any reduction in pain severity was mediated by changes in CME or PAF. Relative to sham, active rTMS applied prior to pain onset led to subsequently lower pain upon chewing and yawning. The neurophysiological results showed that active rTMS increased PAF speed, but did not alter CME. There was no evidence that reductions in pain upon chewing and yawning induced by active rTMS were mediated by changes in PAF or CME. However, regardless of group, faster PAF and higher CME on Day 4 was associated with lower intensity of future pain.

### rTMS delivered prior to a prolonged pain episode protects against future pain severity

Our study showed that 5 days of pre-pain rTMS led to a reduction in future pain upon chewing and yawning within a model of prolonged temporomandibular pain lasting days to weeks. Pain upon functional jaw movement is a key criterion for the diagnosis of clinical TMD [91]. Moreover, previous research has shown that, after an NGF injection to the masseter muscle, pain during chewing and yawning activities is greater compared to other activities, [23;89] which was replicated in the present study. As such, we considered pain upon chewing and yawning as primary outcomes. As anticipated, rTMS showed stronger effects on these measures relative to other outcomes. Further consistent with the idea of jaw movement being the most relevant to the NGF-TMD model, we found stronger reductions in functional limitation of jaw mobility in the active vs. sham group at earlier timepoints. This suggests rTMS delivered prior to the onset of pain can influence both pain itself and pain-related functional impairment.

Taken together, our findings support existing evidence for the analgesic effects of high-frequency rTMS delivered to M1 in clinical populations [1;36;37], and in healthy participants during prolonged pain induced by NGF [13;25;93]. Further, this study is the first to demonstrate a reduction in pain severity when the rTMS intervention is delivered prior to the onset of prolonged pain. Previous studies have shown that single sessions of rTMS delivered prior to pain can subsequently reduce pain sensitivity to transient noxious stimuli relative to sham rTMS or a baseline timepoint prior to rTMS [9;61;69;70;117]. However, the brief nature of these painful stimuli hinder the external validity to clinical pain, which typically lasts for weeks or months [8] . In contrast, the NGF paradigm is one of few experimental pain paradigms that can produce a prolonged pain experience in humans lasting several weeks [8]. Several studies have shown that injecting NGF to the neck, forearm or masseter muscles can mimic symptoms of clinical neck pain [21], lateral epicondylalgia (tennis elbow) [8] and TMD [89] respectively, including the time course (gradual onset and prolonged duration), movement-evoked pain, impaired joint movements, limited function and hyperalgesia (lower pressure pain thresholds).

Given the clinical relevance of the NGF model, our findings have implications for the use of rTMS as a prophylactic pain intervention. A critical limitation of most pain treatments is that they are applied in the chronic stages when maladaptive nervous system plasticity has already occurred and may be difficult to reverse [30;52]. As such, there is growing interest in preventative approaches for the transition from acute to chronic pain. Here, we provide pre-clinical evidence that rTMS delivered prior to a prolonged pain episode may protect against future pain. One application of these findings could be surgical procedures that present a risk of severe post-operative pain and future development of chronic pain [48]. For example, post-operative jaw pain can occur following procedures such as dental implant, orthognathic and maxillofacial surgery, with 5-13% of patients undergoing a dental surgery going on to experience chronic jaw pain [22;79;80;92]. Given a strong predictor of the development of chronic pain after surgery is high pain severity during the acute stage [42;105], the delivery of rTMS prior to surgery could be an effective method to reduce post-operative pain. This would interrupt the transition from acute to chronic pain, whilst also reducing reliance on opioid based treatments that may put individuals at risk of long-term dependence [97]. Future studies are needed to build on these preliminary findings.

### While the mechanisms of rTMS were unclear, PAF and CME predicted future pain

As hypothesised, our findings showed that 5 days of rTMS led to an increase in PAF. This is consistent with previous studies showing an increase in PAF following single sessions of high frequency rTMS [2;67;73], though ours is the first study to demonstrate these effects following a multi-session protocol. Thus, our results suggest that, along with other methods such as exercise [40], visual stimulation [114] and nicotine [66], rTMS can be used as an experimental method of increasing PAF. This has implications for understanding the role of PAF in various other cognitive/sensory processes such as working memory and attention [15;16;41;82]

We did not find an increase in CME following rTMS, which is inconsistent with previous work [25;35]. One possible explanation is that rTMS intensity and location was calibrated relative to the activation of the FDI muscle rather than the masseter muscle. However, a previous study which based the rTMS intensity on the activation of masseter muscle did not show a significant increase in MEP amplitudes following 10Hz rTMS [47]. Another possibility is that effects of rTMS on CME lack reproducibility, as recent research has shown low to moderate reliability of an increases or decreases in CME following high or low frequency rTMS [81]. As such, current rTMS protocols may need further refining to improve the reliability of CME changes following rTMS.

The mediating mechanisms behind the analgesic effects of rTMS remain poorly understood and are seldom investigated [62]. While our study is novel in this regard, we did not find evidence that changes in PAF/CME mediated the effects of rTMS on future pain. This suggests rTMS may have acted on other cortical mechanisms that attenuated future pain. One possibility is other cortical regions (e.g. dlPFC, insular, cingulate cortices) which are *also* activated by M1 rTMS, mediated the prophylactic effects [5;38;44]. Indeed, connectivity between dlPFC and subcortical regions involved in descending pain inhibition has been shown to predict future response to pain interventions [96]. Another possible mechanism is alterations in sensorimotor inhibitory/excitatory processes that cannot be captured by TMS-MEP methodologies, but which can only be captured using combined TMS-EEG methodologies. TMS-EEG allows activity to be measured directly from the cortex rather than from peripheral muscles [18]. We have preliminary evidence showing that GABAergic processes within the sensorimotor cortex (indexed using TMS-EEG) partially mediated the analgesic effects of rTMS during pain [19]. As such, we encourage future studies to determine whether such mechanisms are also involved in the prophylactic effects of rTMS from future pain.

While the present study showed that PAF and CME did not mediate the effect of rTMS on pain, we showed that an individual’s CME or PAF on Day 4, regardless of group allocation, was correlated with future pain intensity. Specifically, lower PAF and CME on Day 4 were associated with higher future pain. This is consistent with previous research showing lower CME during the acute stage of non-specific low back pain predicts higher pain intensity at 6 months follow-up [49], and studies showing that slower PAF prior to experimental pain onset predicts higher future pain [32–34]. Note a relationship between PAF/CME on Day 0 and future pain was not observed. This is likely due to fluctuations in PAF/CME introduced by the rTMS intervention and/or the longer time interval between Day 0 PAF/CME and pain measurements on Day 5-25. Overall, our results suggest that while changes in PAF or CME did not drive the analgesic effects of rTMS, these measures are relevant predictors of future pain sensitivity.

### Strengths and Limitations

This study used a thorough experimental approach with successful blinding of participants to group allocation, roughly equal split of males and females in each group and blinding of the experimenter involved in data collection, pain induction, pre-processing and the statistical analysis plan. One potential limitation of the study is the use of the marked hotspot method for the five sessions of rTMS rather than neuronavigation. The latter has been shown to improve the consistency of coil positioning and orientation across sessions [12]. As such, the use navigation-guided rTMS in the present study may have increased the magnitude of the prophylactic pain effects. Another limitation is that pain catastrophizing and other mood-related measures were not collected following rTMS. This may have provided further information regarding the mediating mechanisms for the protective effect of rTMS, such as changes in affective-emotional dimensions of pain [52].

## Conclusion

Five consecutive days of high frequency rTMS delivered *prior* to pain onset can reduce the intensity of a future episode of prolonged pain, suggesting rTMS may be useful as a prophylactic intervention in some clinical settings, including as an adjunct intervention to a pre-surgical optimisation program aiming to reduce post-surgical pain. Future studies are needed to explore this possibility.

## Acknowledgements

This work was supported by 1R61NS113269-01 from The National Institutes of Health to DAS, SMS, and The Four Borders Foundation to DAS.

